# Photosynthetic living fibers fabrication from algal-bacterial consortia with controlled spatial distribution

**DOI:** 10.1101/2023.05.02.539014

**Authors:** Zitong Sun, Zhengao Di, Yang Zhang, Huilin Wen, Shaobin Zhang, Zhiqian Zhang, Jing Zhang, Ziyi Yu

## Abstract

Living materials that combine active cells and synthetic matrix materials have become a promising research field in recent years. While multicellular systems present exclusive benefits in developing living materials over single-cell systems, creating artificial multicellular systems can be challenging due to the difficulty in controlling the multicellular assemblies and the complexity of cell-to-cell interactions. Here, we propose a co-culture platform capable of isolating and controlling the spatial distribution of algal-bacterial consortia, which can be used to construct photosynthetic living fibers. Through coaxial extrusion-based 3D printing, hydrogel fibers containing bacteria or algae can be deposited into designated structures and further processed into materials with precise geometries. In addition, the photosynthetic living fibers demonstrate a significant synergistic catalytic effect resulting from the immobilization of both bacteria and algae, which effectively optimize sewage treatment for bioremediation purposes. The integration of microbial consortia and 3D printing gives functional living materials that have promising applications in biocatalysis, biosensing, and biomedicine. Our approach provides an optimized solution for constructing efficient multicellular systems and opens a new avenue for the development of advanced materials.

## 1. Introduction

Microbial consortia have shaped the fate of the Earth from the beginning of life. Recent studies of microbial cocultures, in which two or more populations of cells are co-cultured in symbiosis, have inspired a range of living materials that recapitulate the inherent advantages of biological structures[1,2]. Materials are constructed by a variety of microorganisms that grow into functional living materials with diverse physical and chemical properties including self-assembling, self-repairing and the ability to sense environmental stimuli[3,4]. In contrast to monoculture, the use of microbial consortia could create living materials with programmable biochemical properties[5,6,7,8]. By cultivating two or more types of cells within a single system to replicate a natural symbiotic environment, it becomes possible to facilitate the exchange of energy, substances, and signals between different species[9,10,11]. This approach also enables the sharing of labor and metabolic processes, allowing for efficient partitioning of resources[12]. However, spatially controlling the microbial consortia in living materials is not well explored and constructing a co-culture platform that could efficiently distribute and substrates remains challenging[13]. These have greatly hampered our understanding of the emerging properties of microbial consortia and their derived materials.

An ideal microbial consortium should possess an inherent control mechanism that ensures microorganisms are distributed across different spatial domains, thereby preventing fast-growing species from outcompeting slow-growing ones[14]. The control mechanism should also maintain stability and robustness in the production of target metabolites. Immobilization of cells while maintaining their metabolic activity has been demonstrated to have great potential to be a universal co-culturing platform[15]. Typically, microorganisms are selectively encapsulated within a substance, generating a microbe-laden matrix for investigating microbial communication and living materials[16,17]. Hydrogels are commonly used as the matrix for immobilizing microbial consortia due to their biocompatibility, biodegradability, and chemical functionalization capabilities[18]. They allow for the diffusion of small molecules to maintain cell activity and growth while keeping cells in a stable network[19,20,21]. When a viscoelastic and shear-thinning hydrogel is utilized, it enables the bioprinting of cell-laden hydrogels, leading to precise control over the spatial distribution and concentration of microorganisms[22,23]. Although recent research has demonstrated the feasibility of cultivating multiple heterotrophs in hydrogels, the light-driven consortia, which have the potential to enable the reexamination of the global carbon cycle and other biogeochemical processes, have not been thoroughly investigated.

In this study, we report a facile bioprinting tool for the fabrication of photosynthetic living fibers along with the development of the potential bioremediation applications. We utilize a coaxial nozzle to precisely confine algae-bacteria consortia in a biocompatible medium, resulting in the formation of hydrogel fibers that can be further processed into living materials with a well-defined three-dimensional (3D) geometry, facilitating spatial immobilization (Fig. 1A). Interesting, the results demonstrate a noteworthy reduction in methyl orange (MO) coloration through photosynthetic decolorization by a synthetic consortium carefully engineered to distribute metabolic tasks between *Chlorella vulgaris* (*C. vulgaris*) and *Bacillus subtilis* (*B. subtilis*) (Fig. 1B). The results show that the efficiency of MO decolorization in algae-bacteria consortiums is dependent on their spatial distribution, highlighting the crucial role of mass transfer processes in determining the efficacy of photosynthetic living fibers. Taking together, the integration of algae-bacteria consortia into hydrogel fibers through 3D printing creates an ideal platform for advancing both materials engineering and microbial algal-bacterial associations[24]. To fully realize the potential of these materials, it is essential to improve the accuracy of models and enhance our ability to manipulate cellular behavior in the future.

**Fig. 1.**
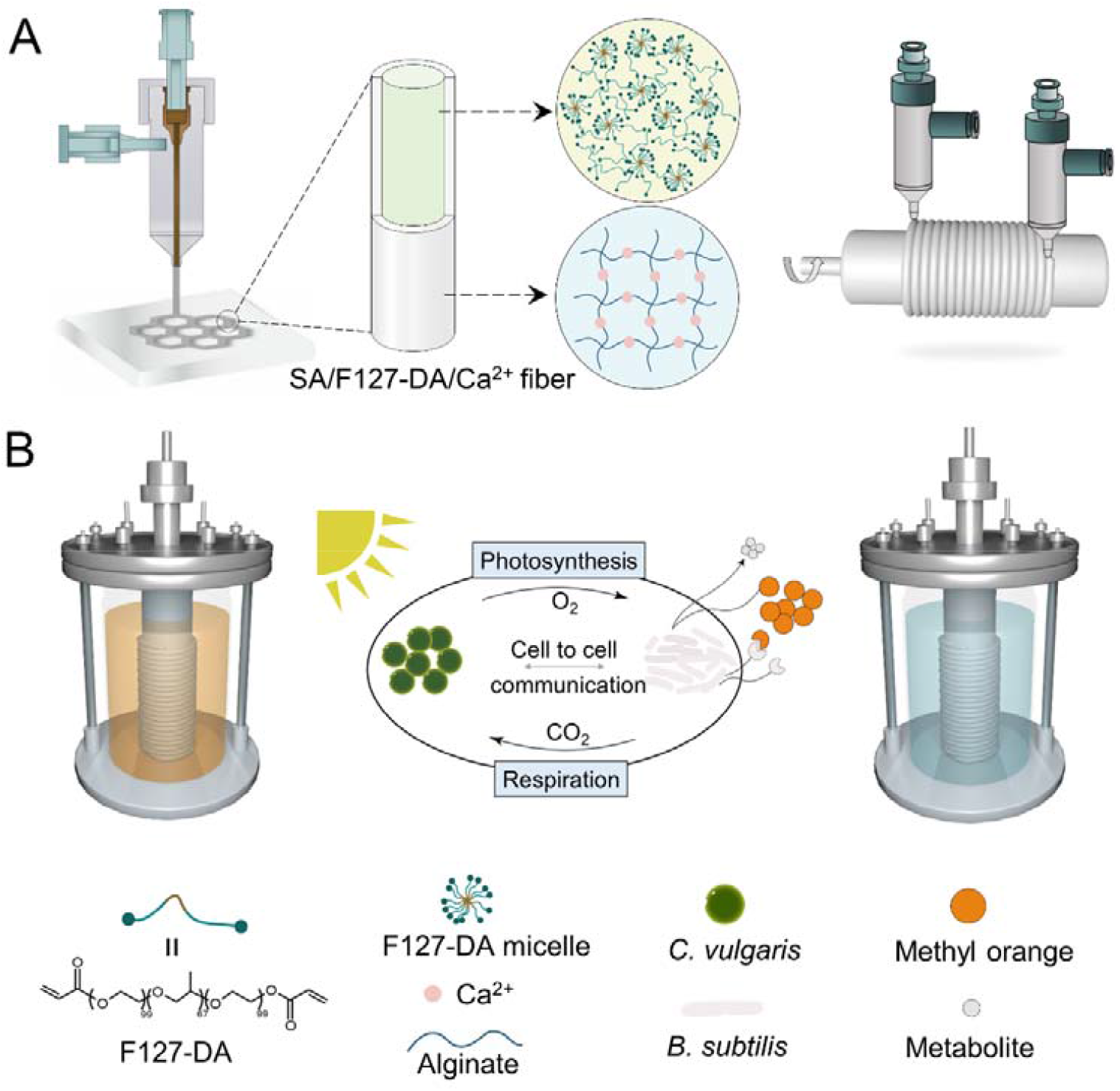
Overview of coaxial extrusion fabrication of microbial reactors based on bacteria-algae symbiosis. (A) Schematic illustration of the preparation of hydrogel fibers by coaxial extrusion. F127-DA solution was rapidly formed as a hydrogel network by free radical polymerization reaction under blue light, and SA solution was rapidly cross-linked with Ca^2+^. (B) Schematic illustration of microbial reactor based on bacteria-algae symbiosis. Microbial reactor could be used for MO decolorization under the joint action of *B. subtilis* and *C. vulgaris*.

## 2. Materials and methods

### 2.1. Materials

BG11 medium was purchased from Qingdao Hi-Tech Industrial ParkHaibo Biotechnology Co., Ltd. All other reagents (sodium alginate (SA), calcium chloride (CaCl_2_), ammonium chloride (NH_4_Cl), potassium nitrate (KNO_3_), sodium nitrite (NaNO_2_), monopotassium phosphite (KH_2_PO_3_), methyl orange (MO) lithium phenyl (2,4,6-trimethylbenzoyl) phosphinate (LAP), sodium chloride (NaCl), tryptone, yeast extract, d(+)-sucrose (Suc), Sodium acetate (Ace), glucose (Glu), lactose (Lac), sodium hydroxide (NaOH), hydrochloric acid (HCl), polyether F127 (F127) chloroform (CHCl_3_) triethylamine (C_6_H_15_N), acryloyl chloride (C_3_H_3_ClO))were purchased from Aladdin Shanghai Reagent Co., Ltd (Shanghai, China). All aqueous solutions were deionized water (resistivity of 18.2 MΩ·cm at 25 °C) treated with the Milli-Q^™^ reagent system.

### 2.2. Microorganisms and culture conditions

*Chlorella vulgaris* (GY-D19 Chlorella sp.) was obtained from the Shanghai Guangyu Biological Technology Co., Ltd. *C. vulgaris* was cultivated in flasks containing 200 mL BG11 medium at 25 °C for 7 days with a 12 /12 h light-dark cycle at a light intensity of 6000 lux.

*Bacillus subtilis* (*B. subtilis*) was obtained from Professor Jiang Min’s group at Nanjing Technology University. GFP expressing *Bacillus subtilis* (GFP-*B. subtilis*) was obtained from Hangzhou Baosai Biotechnology Co., Ltd. *B. subtilis* was inoculated in 5 mL of LB medium and incubated in a shaker at 35 °C, 200 rpm for 8 h. The LB medium contained 10 g/L tryptone, 5 g/L yeast extract, and 10 g/L NaCl.

The quantification of *C. vulgaris* was performed by the hemocytometer method using an optical microscope. The total number of *B. subtilis* was counted by the plate count method. GFP-*B. subtilis* was utilized to study the distribution of *B. subtilis* within core-shell fibers and to assess its proliferation in co-culture, but not to conduct MO decolorization experiments.

### 2.3. Synthesis of F127-DA

First, polyether F127 (90 mmol, 10 g) was added to a 250 mL flask and dried in a vacuum drying chamber at 80 °C for 4 h. After F127 cooled to room temperature, CHCl_3_ (0.99 mmol, 0.119 g) was added to the flask to dissolve F127 in an ice bath. C_6_H_15_N (7.87 mmol, 0.7968 g) was then added and stirred until well mixed. To a constant pressure burette containing 20 mL of CHCl_3_ (0.249 mmol, 0.0298 g), add a C_3_H_3_ClO (9.55 mmol, 0.11 g) in slow drops to the flask. Then, the reaction was carried out at room temperature for 48h, and the precipitate was filtered off. After spin evaporation, the filtrate was added to an excess of C_4_H_10_O, and a white precipitate was obtained by filtration. The precipitate was dissolved in a small amount of CHCl_3_ and then filtered after precipitation in C_4_H_10_O, and the resulting white solid was dried in a vacuum drying chamber. The synthesis principle and ^1^H NMR spectrum of F127-DA (Fig. S1).

### 2.4. Bio-Ink preparation

Centrifuge *C. vulgaris* (OD_680_=1.0) and *B. subtilis* (OD_600_=0.9) at 4500 rpm for 5 min, then mix the resulting microbial sediment with SA solution to embed the microorganisms in the shell phase. The core phase flow is a 20wt% F127-DA solution containing excess Ca^2+^. In subsequent experiments, microorganisms in the shell phase will be immobilized with the SA solution, while microorganisms in the core phase will be immobilized with the 20wt% F127-DA solution containing excess Ca^2+^.

### 2.5. Printability of the material

3D printing of the biomaterials was performed by a commercial bioprinter (EFL-BP-6601, Yongqinquan Intelligent Equipment Co., Ltd., China) with the following printing process: F127-DA/Ca^2+^ and SA were transferred into a sterile syringe and fixed on a coaxial printing nozzle. The printing settings were: extrusion speed of 0.2 mL/min and 0.5 mL/min for the core and shell phases, respectively, print head speed of 360 mm/min, and dispensing needle size of (0.42×0.72 mm). The print substrate was placed on a 35 °C printing platform and the prepared hydrogel fibers were cross-linked under blue light (405 nm) and then transferred to 0.5 M CaCl_2_ solution (pH = 7) for 30 min. Finally, the scaffolds were washed with deionized water.

### 2.6. Cell viability

Printed *C. vulgaris*-laden scaffolds were incubated for 14 days at 25 °C with a light intensity of 6000 lux in a 12 /12 h light/dark cycle. After 0, 7, or 14 days, samples were taken for observation. The growth of the scaffolds was observed by visual and fluorescence microscope (IX73, Olympus Corporation, Japan). At the same time, the proliferation of the same amount of microbial consortia in co-cultivation and single culture (in a free state) was compared. *C. vulgaris* was imaged using a fluorescence microscope with excitation/emission wavelengths of 480/628 nm and GFP-*B*.*subtilis* at excitation/emission wavelengths of 470/535 nm.

### 2.7. Controlled fabrication of fibers

The size of the inner diameter of the fibers was controlled by adjusting the flow rate of the core phase and the diameter of the needle inside the chip. The flow rate of the core phase was set to 100, 200, 300, 400, and 500 μL/min, and the prepared fibers were observed under a microscope. The inner needle diameter of the chip was adjusted to 0.42, 0.61, 0.72, 1.11, and 1.43 mm and the prepared fibers were observed under the microscope. The experimental procedure maintains a single variable.

### 2.8. Decolorization of MO by *B. subtilis*/*C. vulgaris*-laden fibers

Synthetic wastewater: The synthetic wastewater was used in this study and the compositions were 400 mg/L glucose, 22.16 mg/L NH_4_Cl, 6.57 mg/L KNO_3_, 1.08 mg/L NaNO_2_, 5.09 mg/L KH_2_PO_3_ and 10mg/L MO [25].

The decolorization experiments were divided into three groups. The *B. subtilis/C. vulgaris*-laden fibers were inoculated in serum bottles containing 10 mL of wastewater and decolorized in a shaker at 35 °C. After purging with Argon to ensure anaerobic conditions, the serum bottles were sealed with rubber stoppers. Samples taken every 12 hours were centrifuged at 5000 rpm for 5 min to isolate any microorganisms that might have leaked, and the decolorization rate of the dye at maximum absorbance was determined by UV spectrophotometry. The decolorization percent was computed by the following formula[26,27].

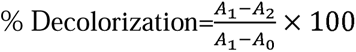

where A_0_ is the O.D. of deionized water, A_1_ is the initial O.D. of the sample, and A_2_ is the final O.D. of the sample.

### 2.9. Optimization of the decolorization conditions of MO by photosynthetic living fibers

#### 2.9.1. Effect of spatial distribution on the decolorization process

*B. subtilis* and *C. vulgaris* can be encapsulated in the core and shell phases of fibers, respectively. Four different types of encapsulation forms are discussed in this paper. *B. subtilis*/*C. vulgaris* co-culture system 1 (shell phase: *B. subtilis*, core phase: *C. vulgaris*), co-culture system 2 (shell phase: *C. vulgaris*, core phase: *B. subtilis*), co-culture system 3 (shell phase: *B. subtilis*/*C. vulgaris*), co-culture system 4 (core phase: *B. subtilis*/*C. vulgaris*). Taking co-cultivation system 1 as an example, *B. subtilis* was mixed with shell phase SA solution after centrifugation, while *C. vulgaris* was mixed with core phase F127-DA/Ca^2+^ following centrifugation, and core-shell fibers were prepared through a coaxial extrusion device. We used a similar method to prepare the other three *B. subtilis*/*C. vulgaris*-laden fibers. Decolorization experiments were performed using the four co-culture systems described above. *B. subtilis*-laden monoculture hydrogel served as a control. Decolorization experiments were performed by equal amounts of free cells under the same decolorization conditions as for immobilized cells.

#### 2.9.2. Effect of the ratio of *B. subtilis* and *C. vulgaris* on the decolorization process

The experiment aimed to explore the effect of the *B. subtilis* and *C. vulgaris* ratio on the decolorization process. The ratio of *B. subtilis*: *C. vulgaris* was varied as 1:0, 2:1, 1:1, 1:2, and 1:3, and the resulting mixtures were embedded in hydrogel fibers for decolorization experiments. The experiment maintains a single variable. Wastewater samples were taken every 12 hours and analyzed by UV spectrophotometry.

#### 2.9.3. Effect of carbon source on the decolorization process

The experiment aimed to investigate the effect of different carbon sources on the decolorization process. Five carbon sources were used: carbon-free (NC), sucrose (Suc), sodium acetate (Ace), lactose (Lac), and glucose (Glu). Decolorization experiments were performed using *B. subtilis/C. vulgaris*–laden fibers under different carbon sources. The experiment maintains a single variable. Wastewater samples were taken every 12 hours and analyzed by UV spectrophotometry.

#### 2.9.4. Effect of pH on the decolorization process

Experimental water samples were adjusted to pH levels of 4, 5, 6, 7, and 8 using 1M HCl or 1M NaOH. MO decolorization experiments were then performed using *B. subtilis*/*C. vulgaris*-laden fibers at each pH level. The experiment maintains a single variable. Wastewater samples were taken every 12 hours and analyzed by UV spectrophotometry.

### 2.10. Reuse and leakage

In the reuse experiment, *B. subtilis*/*C. vulgaris*-laden fibers were inoculated into fresh MO wastewater for cyclic decolorization experiments without adding additional *B. subtilis* and *C. vulgaris*. The decolorization time for each cycle was limited to 48 h, and wastewater samples were taken every 12 hours and analyzed by UV spectrophotometry. Encapsulated *B. subtilis* and *C. vulgaris* within hydrogel fibers may experience proliferation during the cyclic experiment. On the 4 and 8 days, we took 100 μL of wastewater samples and counted the leaked bacteria using the plate counting method. On the 8 day of incubation, we took 1 mL of the wastewater sample, centrifuged it at 5000 rpm for 5 min, and used the microbe precipitate for MO decolorization experiments.

### 2.11. Analysis of decolorization products

#### 2.11.1. UV-Vis spectral analysis

To track dye decolorization and metabolite formation, we used a UV-Vis spectrophotometer (UV-3600, SHIMADZU, Japan) to analyze the variation in UV-Vis spectra (ranging from 200 nm to 800 nm).

#### 2.11.2. HPLC analysis

We used HPLC (Agilent, LC1260 Infinity‖) equipped with a C-18 column and UV-Vis detector to analyze the azo bond breakage products of MO. The eluent was a mixture of ammonium acetate (10 mM, pH 4) and methanol in a 50:50 volume ratio, with a flow rate of 0.5 mL/min[26]. We filtered the samples through a 0.22 μm filter tip and detected metabolites at 245 nm.

## 3. Results and discussion

Through a coaxial extrusion device, 2wt% SA solution focused the core phase 20wt% F127-DA solution with Ca^2+^, F127-DA rapidly formed a hydrogel network by free radical polymerization reaction under blue light initiator, and SA was rapidly cross-linked with Ca^2+^ (Fig. 1A, 2A). The SA/F127-DA/Ca^2+^ hydrogel fibers prepared by ionic crosslinking and photo-crosslinking had a core-shell structure that were morphologically continuous and homogeneous, and also had certain mechanical properties (Fig. 2B, 2C). The freeze-dried SA/F127-DA/Ca^2+^ fibers showed a separation of core and shell (Fig. 2D). The SEM image of calcium alginate formed after SA cross-linked with Ca^2+^ (Fig. 2G), calcium alginate presented a unique sheet-like structure. The SEM image of F127-DA injected into the core phase (Fig. 2H), the structure of F127-DA was tight with minimal pores.

**Fig. 2.**
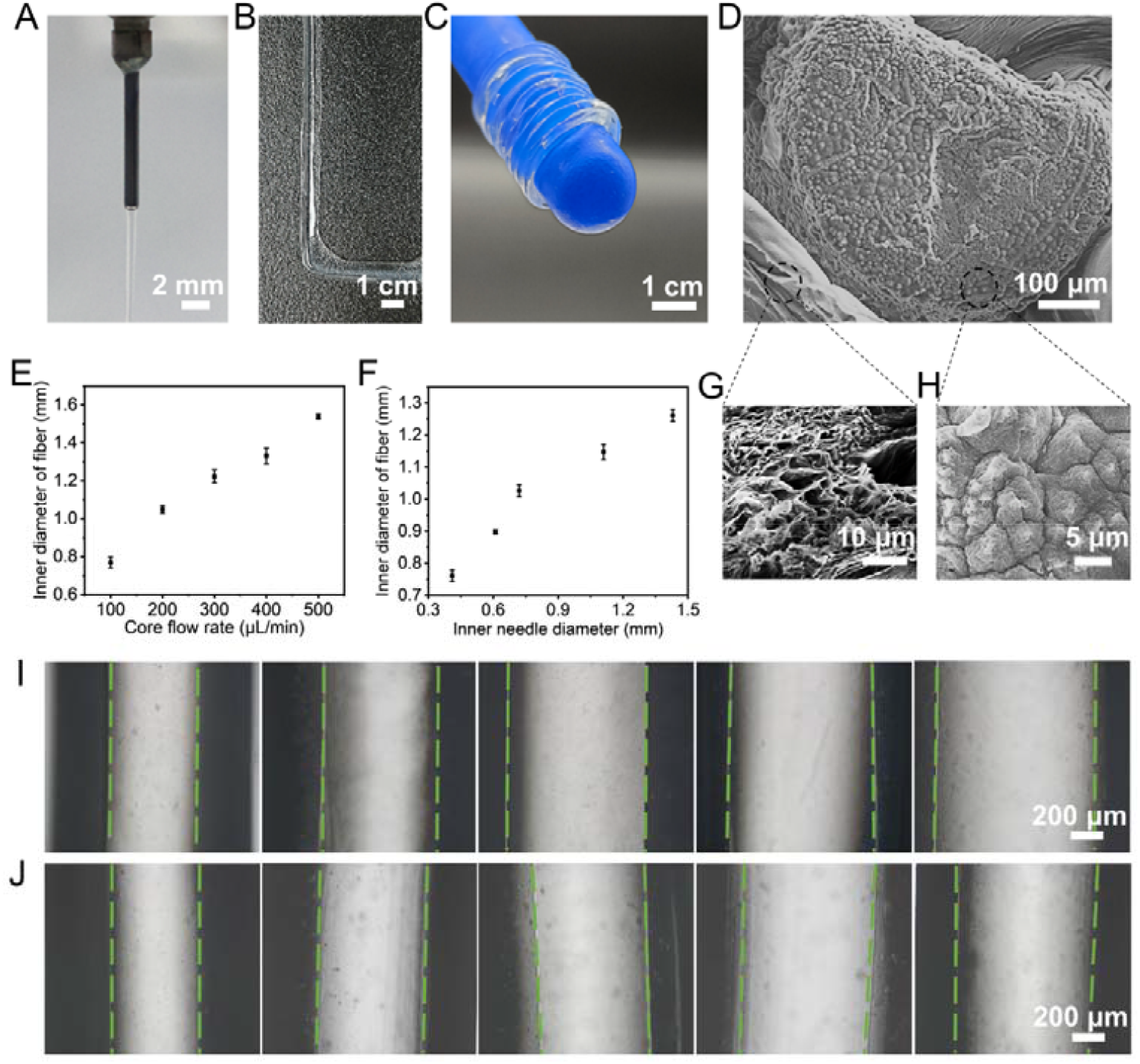
Controlled fabrication of SA/F127-DA/Ca^2+^ core-shell fibers. (A) Image of fabrication of core-shell fibers using a coaxial extrusion device. (B-C) Images of SA/F127-DA/Ca^2+^ hydrogel fibers. (D) SEM images of SA/F127-DA/Ca^2+^ core-shell fibers. (G), (H) Close-up SEM images of the shell phase SA (G) and core phase F127-DA (H). (E) The relationship between fiber inner diameter and core flow rate. The core flow rates were 100, 200, 300, 400 and 500 μL/min, respectively, and (I) shows microscope images corresponding to each flow rate. (F) The relationship between fiber inner diameter and inner needle diameter. Inner needle diameters were 0.42, 0.61, 0.72, 1.11 and 1.43 mm, respectively, and (J) shows microscope images corresponding to each inner needle diameter.

In the process of 3D bioprinting, the control of fiber size is very important for constructing complex structures. In this experiment, we controlled fiber sizes by varying the flow rate of the core phase solution and the inner needle diameter. When the core flow rate was increased, the internal diameter of the fiber increased (Fig. 2E) and the core space became larger (Fig. 2I). This was due to a decrease in the effective diffusivity of core phase Ca^2+^ when the core flow rate increased, resulting in an increase in the internal diameter of the fiber[28]. Similarly, an increase in the inner needle diameter also led to a larger fiber internal diameter (Fig. 2F, 2J). This may be due to the fact that as the diameter of the inner needle gradually increased, the concentration of Ca^2+^ gradually decreased[29]. The fiber size in subsequent experiments was kept at 800 μm to allow enough space for cell proliferation and control the same level of nutrient diffusion.

To assess the printability of the biomaterials, two pre-defined geometric patterns were printed and visually inspected at 0, 7, and 14 days of light-mixed growth (Fig. 3A, 3B). Chlorophyll auto-fluorescence was used to observe the growth of *C. vulgaris* under a fluorescence microscope at 0, 7, and 14 days of incubation (Fig. 3C). During the bioprinting and culturing process, *C. vulgaris* was encapsulated within the hydrogel matrix and the SA/F127-DA/Ca^2+^ hydrogel fibers were proved suitable for cell growth while maintaining the 3D structure. In addition to the monolayer structure, we also tried to create a multilayer complex structure by depositing bioink layer by layer on top of the previously printed one (Fig. 3D). During the incubation process, the hydrogel fibers deposited layer by layer can maintain their regular structure without deformation (Fig. S2). These experimental results demonstrate the ability of SA/F127-DA/Ca^2+^ bioinks to build 3D structures as well as complex patterns. It can be used as a living material and scaffold for biomedicine and tissue engineering[30,31,32].

**Fig.3.**
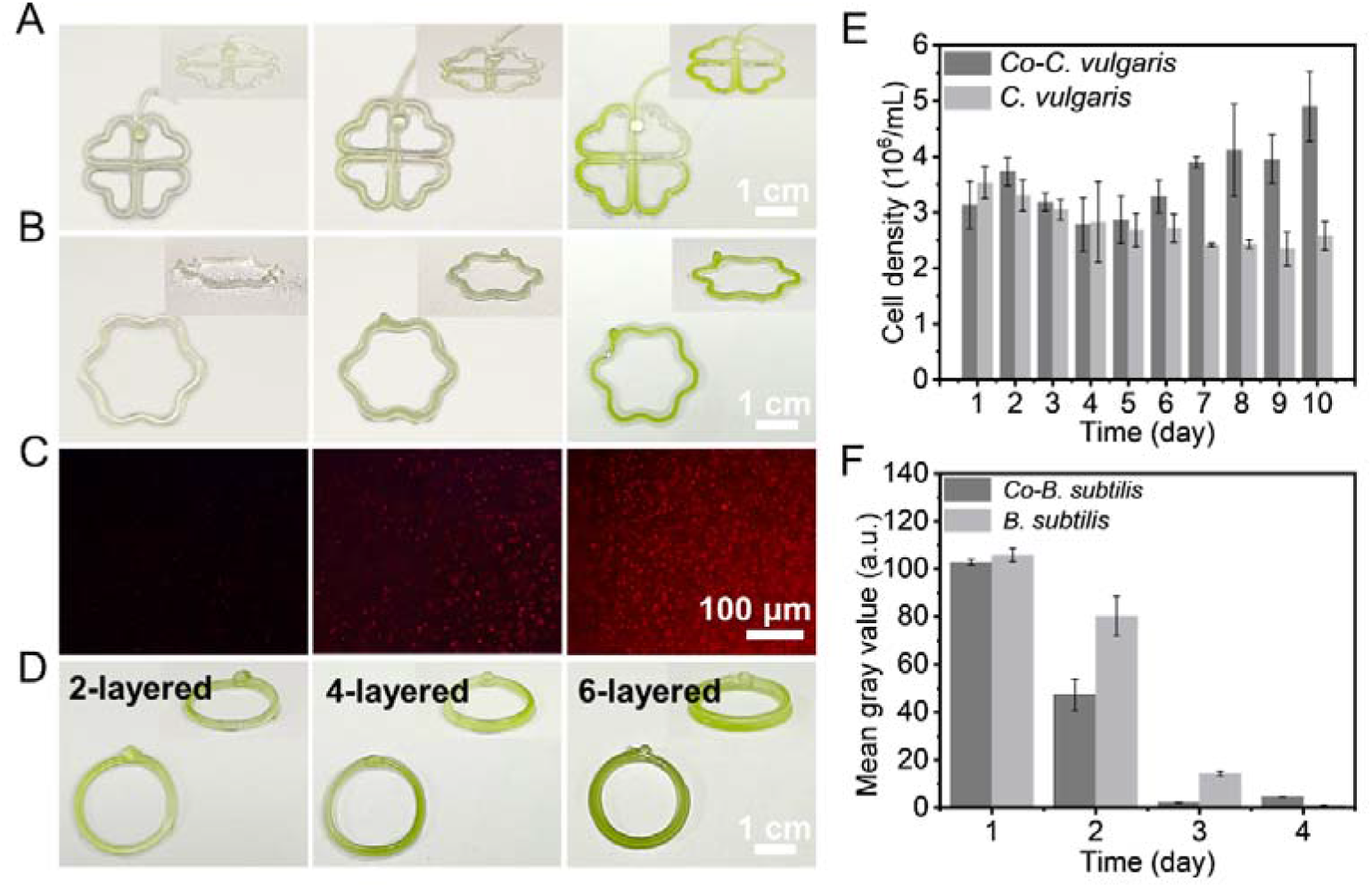
SA/F127-DA/Ca^2+^ photosynthetic living fibers fabricated by 3D bioprinting. (A-B) Different shapes of living fiber were fabricated by 3D bioprinting, and the proliferation of cells was observed at 0, 7 and 14 days after printing. (C) Proliferation of *C. vulgaris* was observed under a fluorescence microscope at 0, 7 and 14 days after printing. (D) Side and top views of circular patterns prepared by layer-by-layer deposition. (E) Cell density of *C. vulgaris* in synthetic wastewater during the monoculture and the co-culture of *B. subtilis*/*C. vulgaris*. (F) Proliferation of GFP-*B. subtilis* in synthetic wastewater during GFP-*B. subtilis*/*C. vulgaris* co-culture evaluated by fluorescence intensity.

The proliferation of *B. subtilis* and *C. vulgaris* co-cultured in synthetic wastewater was evaluated using a cell counting method and fluorescence intensity analysis. The results showed that the proliferation rate of *C. vulgaris* in the co-culture was higher than that in monoculture after 5 days (Fig. 3E), whereas the fluorescence intensity of co-cultured GFP*-B. subtilis* was weaker compared to its monoculture (Fig. 3F).

The SA/F127-DA/Ca^2+^ hydrogel system will provide a platform that can encapsulate microorganisms in different spaces using the core and shell phases of the hydrogel fibers to construct spatially separated microbial communities. This study explored four different encapsulation methods for distributing *B. subtilis* and *C. vulgaris* in different spaces of the core-shell fiber (Fig. 4A), as described in Section 2.9.1. Fluorescence microscopy images of these four encapsulation methods (Fig. 4B).

**Fig. 4.**
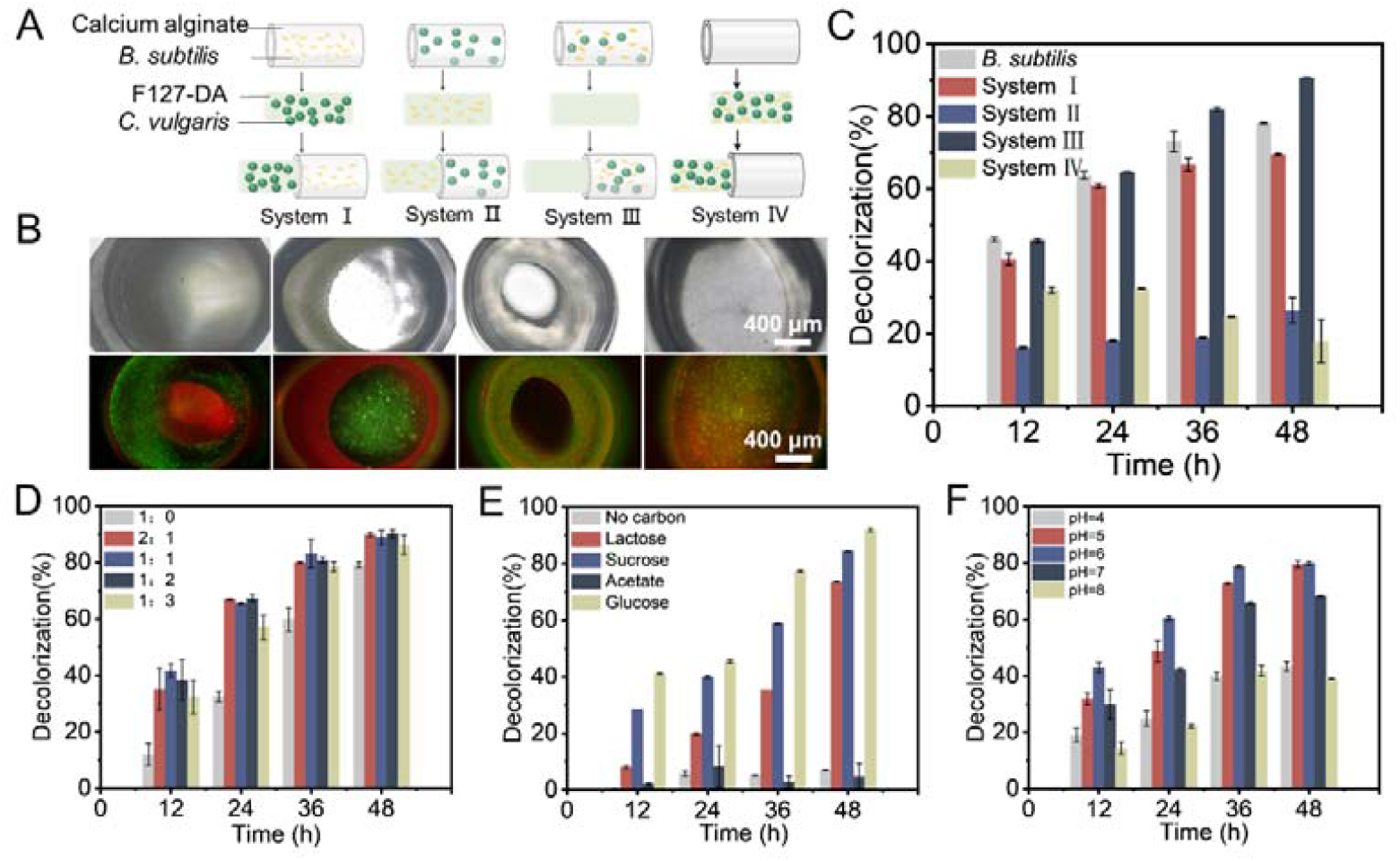
*B. subtilis*/*C. vulgaris*-laden fiber for MO decolorization and optimization of conditions. (A) Four encapsulated forms of GFP-*B. subtilis* and *C. vulgaris* with different spatial distributions. (B) Fluorescence microscope images of four encapsulation forms. (C) MO decolorization conducted by the four encapsulation forms of *B. subtilis*/*C. vulgaris*-laden fibers. (D) Effect of the ratio of *B. subtilis* and *C. vulgaris* on the decolorization process. (E) Effect of carbon source on the decolorization process. (F) Effect of pH on the decolorization process.

To investigate how the spatial distribution of microbial consortia affects their biological activities, we tested *B. subtilis*/*C. vulgaris*-laden fibers in different encapsulated forms for MO decolorization (Fig. 1B). The results indicated that the decolorization rate of co-cultured system 3 was up to 90%, which was higher than that of the monoculture *B. subtilis*-laden hydrogel (Fig. 4C). The lowest decolorization efficiency of co-cultured system 4 might be due to the fact that *B. subtilis* and *C. vulgaris* were encapsulated in the core phase of the hydrogel undergoing a more complex diffusion process. Comparison of the MO decolorization rate of the co-cultured system 3 with the free state of *B. subtilis*/*C. vulgaris* (Fig. S3) showed that the free state of *B. subtilis*/*C. vulgaris* was more efficient in decolorization. This difference may be due to variations in intraparticle diffusion between the free and fixed systems and a higher dispersion of the free microorganisms[33].

The UV-Vis spectra of MO exhibited a single absorbance peak at 463 nm. As the decolorization time progressed, the peak at 463 nm diminished and a new peak appeared in the UV region, indicating the emergence of smaller molecular weight metabolites. The disappearance of the 463 nm peak indicated the breakdown of the N=N bond (Fig. S4). HPLC analysis (Fig. S4) was conducted on the decolorized wastewater samples of *B. subtilis* and *B. subtilis*/*C. vulgaris*. The data indicated that *B. subtilis*/*C. vulgaris* exhibited superior by-product removal efficiency compared to *B. subtilis*. We hypothesize that *B. subtilis* facilitates the reductive cleavage of the azo bond in MO, while *C. vulgaris* absorbs and metabolizes the resulting aromatic amines in the subsequent decolorization process[34,35,36].

We investigated the effects of *B. subtilis*/*C. vulgaris* ratio, different pH, and carbon sources on the decolorization rate of MO (Fig. 4D-F). The results showed that there was little difference in the decolorization rate of MO when different ratios of *B. subtilis*/ *C. vulgaris* were applied (Fig. 4D), and the decolorization efficiency of the *B. subtilis*/*C. vulgaris*-laden fiber was better than *B. subtilis*-laden fiber. However, the *C. vulgaris*-laden fiber did not exhibit the ability to decolorize MO when compared to the blank fibers (Fig. S5). The addition of carbon source significantly promoted decolorization, and glucose as a carbon source had the highest decolorization rate but short-chain fatty acids sodium acetate had less effect on the decolorization rate (Fig. 4E). We believe that pH also affects the decolorization rate and *B. subtilis*/*C. vulgaris* showed the best decolorization in a slightly acidic environment with pH 6 (Fig. 4F).

We also found that *B. subtilis*/*C. vulgaris*-laden fibers were more resistant and showed better adapted to unfavorable conditions than *B. subtilis*-laden fibers (Fig. S6). This phenomenon may be caused by the complex interactions between bacteria and algae[37,38]. The interdependence of bacteria and algae promotes faster growth, robustness to environmental oscillations, and minimized invasion by other species[39]. Furthermore, microalgae can serve as a secondary habitat for bacteria to protect them from harsh environments[39,40].

The reusability of microbe-laden hydrogel is also an important consideration for the application of photosynthetic living fibers in bioremediation. We applied the four previously described *B. subtilis*/*C. vulgaris* fibers with different spatial distributions to MO cycle decolorization (Fig. 4A) to evaluate the reusability and encapsulation ability of the hydrogel during continuous decolorization. Over 8 days, symbiotic systems 2 and 4 exhibited progressively faster decolorization rates, possibly due to the proliferation of *B. subtilis* inside the fibers into the shell phase (Fig. 5A). Meanwhile, symbiotic system 3 consistently showed the fastest decolorization rate among all systems tested, including systems 1, 2, and 4 (Fig. 5A,5B). The decolorization rate of symbiotic system 3 gradually decreased over four cycles, with the fourth cycle achieving 43% within 48 hours (Fig. 5B). The subsequent decolorization experiment using the same fiber showed only a slight improvement, with the rate increasing to 74% on the 6 day (Fig. 5C).

**Fig. 5.**
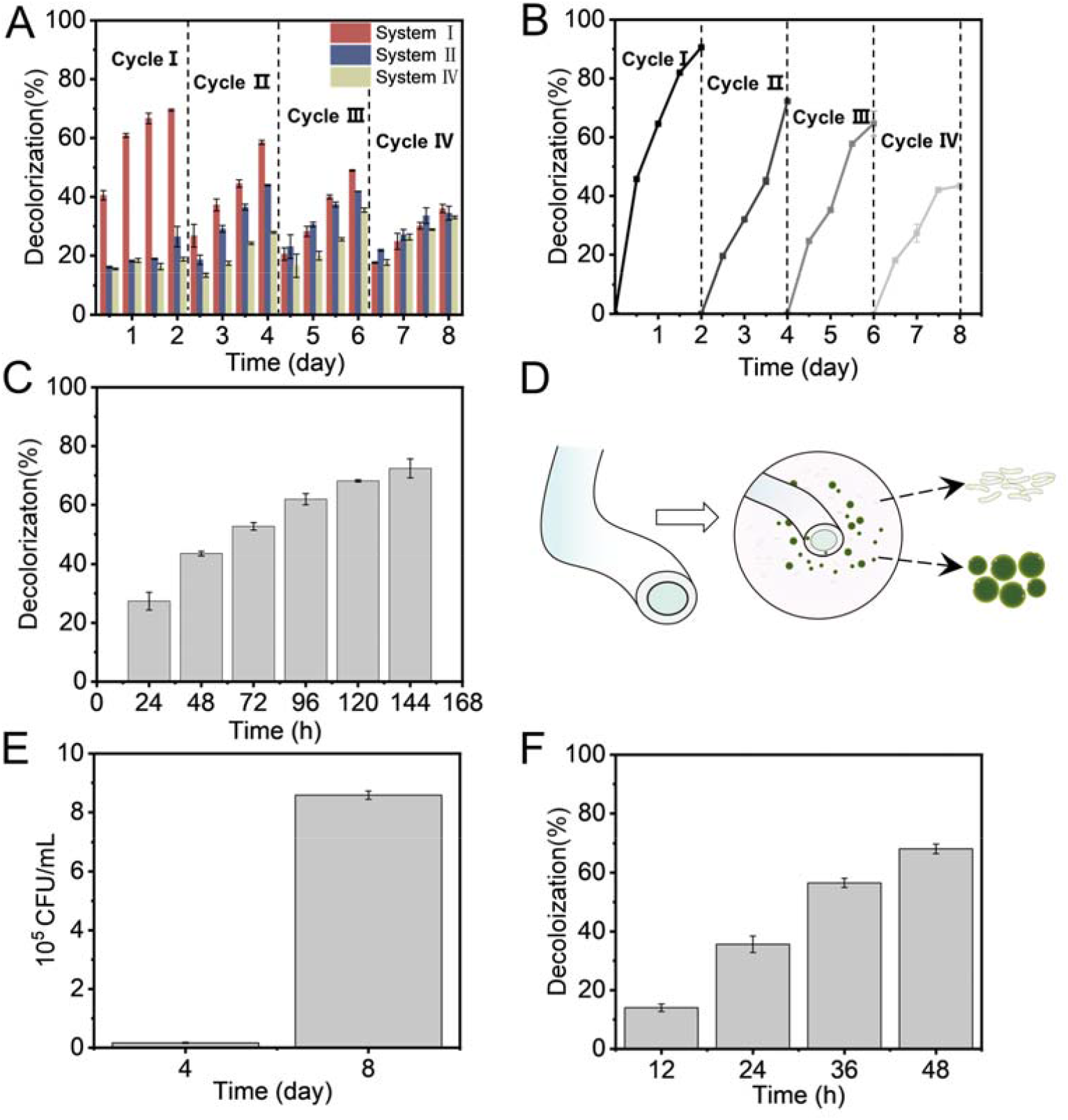
The reusability and encapsulation ability of *B. subtilis*/*C. vulgaris*-laden fiber. (A) Application of *B. subtilis*/*C. vulgaris* Co-culture in to forms of encapsulation system 1,2,4 to MO cyclic decolorization. (B) Application of *B. subtilis*/*C. vulgaris* Co-culture in to form of encapsulation system 3 to MO cyclic decolorization. (C) After four cycles of decolorization, co-culture system 3 continued to undertake MO decolorization. (D) Leakage of *B. subtilis* and *C. vulgaris* immobilized in hydrogel fibers. (E) Leakage of microorganisms on the 4 and 8 days of the decolorization process. (F) MO decolorization process of leaked microbial consortia within 48h.

During the 8-day decolorization process, the decolorized MO wastewater gradually became turbid, which was attributed to the microorganisms escaping from the microbe-laden fibers (Fig. 5D,5E). When the leaked microorganism from the immobilized substrate was applied for decolorization, the rate of decolorization reached 68% for 48 h under the same experimental conditions, which proved that the leaked microorganism contained *B. subtilis* (Fig. 5F). Additionally, our study revealed that microorganisms immobilized in SA/F127-DA/Ca^2+^ fibers not only leaked into the surrounding solution but also proliferated towards the core phase (Fig. S7). Notably, the core-shell fibers we prepared do not entirely isolate microorganisms, and microorganisms immobilized in different spaces may grow into the same space during actual incubation.

## 4. Conclusion

In summary, we have demonstrated a type of photosynthetic living fibers prepared through 3D bioprinting technology to enable the controlled spatial fixation of algae-bacteria consortia in a biocompatible hydrogel medium. This approach allows for the creation of complex 3D structures and ideal cell spatial distribution. The microbe-laden fibers exhibit excellent catalytic properties, making them effective for environmental remediation. *B. subtilis*/*C. vulgaris* with different spatial distributions exhibit different growth and metabolic activities when applied to synthetic wastewater treatment. These findings provide a foundation for developing living materials that support complex multicellular systems. Future research should focus on enhancing our ability to manipulate cellular behavior and improve model accuracy, enabling these materials to realize their full potential in diverse fields such as energy harvesting, biomedical devices, biosensors, and biocatalytic devices.

## Supporting information

This file serves as a supplement to the manuscript data

## Ethics statement

The manuscript does not include any clinical trials. No ethics approval and consent to participate are required for this manuscript.

## Data availability

The raw and processed data required to reproduce these findings are available from the corresponding authors upon request.

## Acknowledgments

This work was supported by National Key Research and Development Program of China (2021YFC2104300) and the National Natural Science Foundation of China (52003119, 52273207). The authors also would like to acknowledge the support of Natural Science Foundation of Jiangsu Province (BK20221314) and State Key Laboratory of Materials-Oriented Chemical Engineering (SKL-MCE-22A06, KL20-02).

