## Supplementary material for "Photosynthetic living fibers fabrication from algal-bacterial consortia with controlled spatial distribution": This file serves as a supplement to the manuscript data

1. Supporting Figures

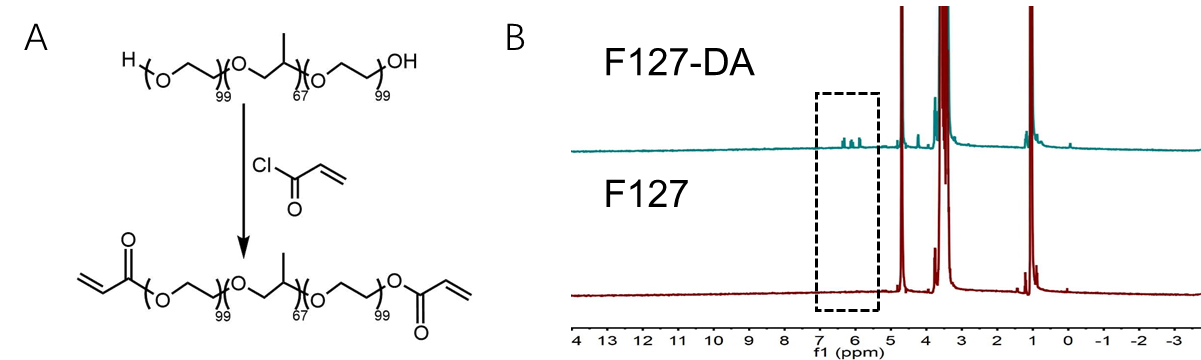

**Fig. S1.** **Synthesis of F127-DA.** (A) Synthesis principle of F127-DA. (B) ^1^H NMR spectrum of F127-DA.

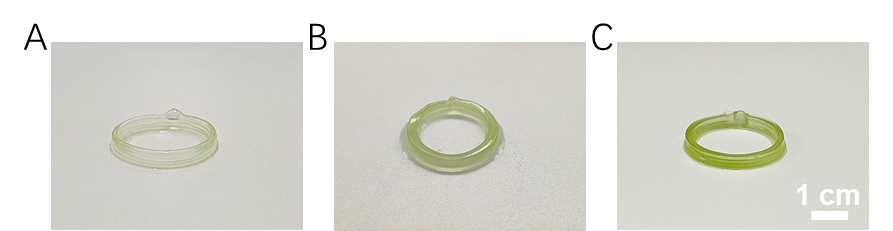

**Fig. S2.** **Incubation of the multilayer scaffold observed on days 0 (A), 7 (B), and 14(C), respectively.**

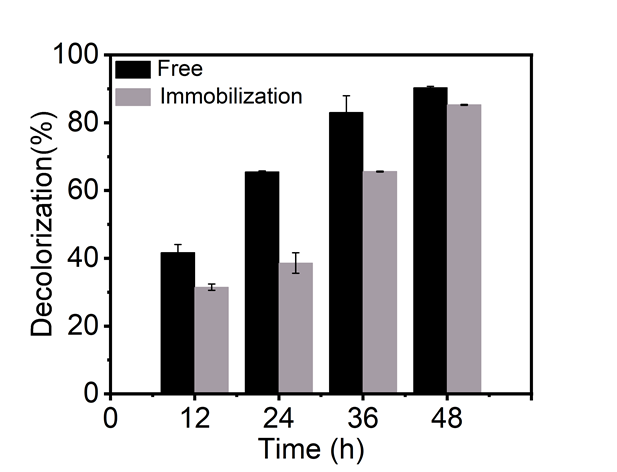

**Fig. S3. Comparison of MO decolorization rates between free and immobilized *B. subtilis/C. vulgaris* co-culture.**

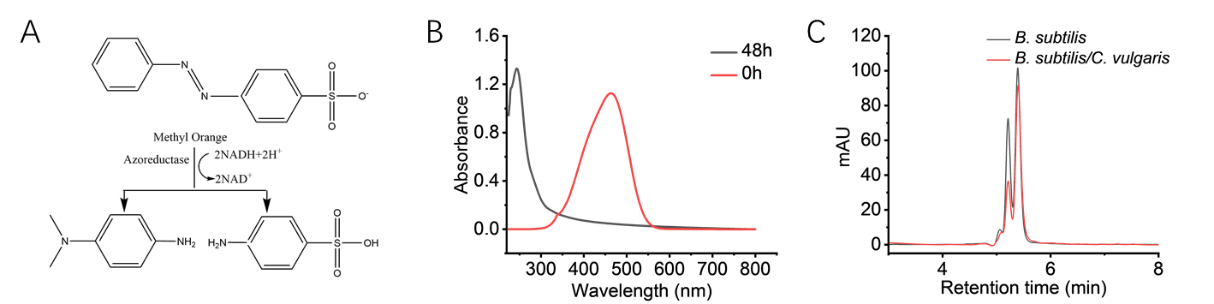

**Fig. S4. MO degradation products and analysis.** (A) Principle of MO degradation. (B) UV-Vis spectra of MO before and after decolorization by *B. subtilis*/*C. vulgaris*. (C) HPLC analysis of *B. subtilis* and *B. subtilis*/*C. vulgaris* decolorized wastewater.

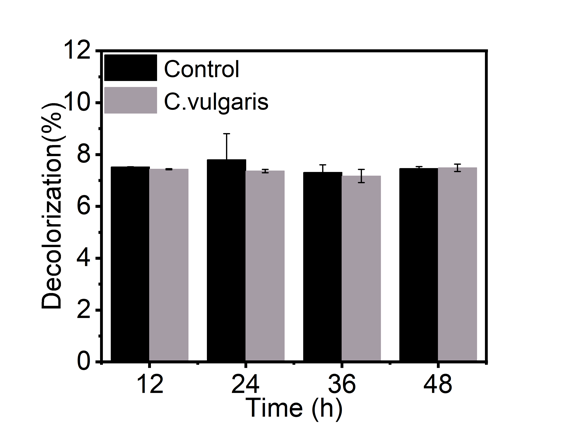

**Fig. S5. Decolorization rates of *C. vulgaris*-laden fiber and blank fiber.**

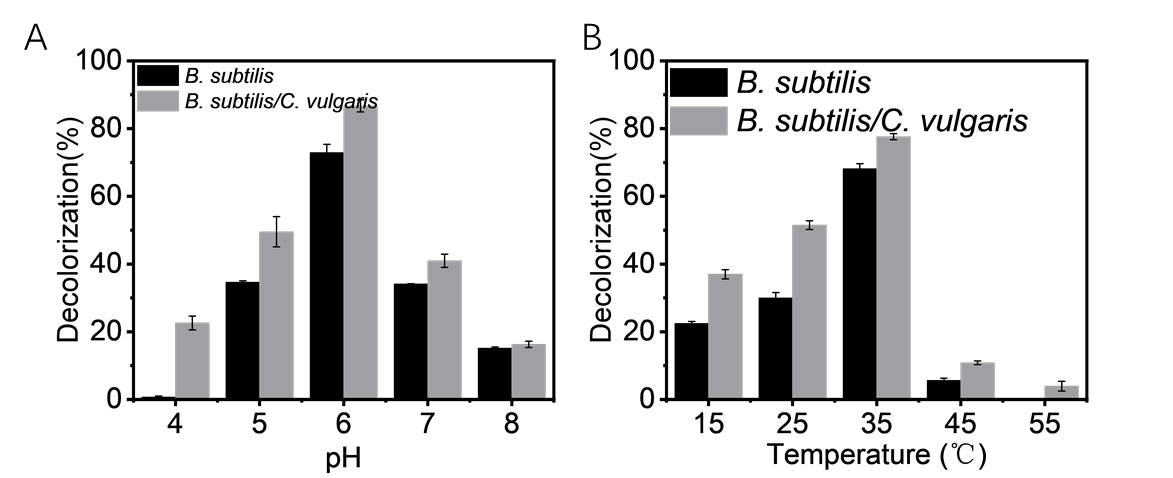

**Fig. S6. Decolorization process of MO under different conditions.** (A) Decolorization rates of *B. subtilis*/*C. vulgaris*-laden fiber and *B. subtilis*-laden fiber under different pH. (B) Decolorization rates of *B. subtilis/C. vulgaris*-laden fiber and *B. subtilis*-laden fiber under different temperatures.

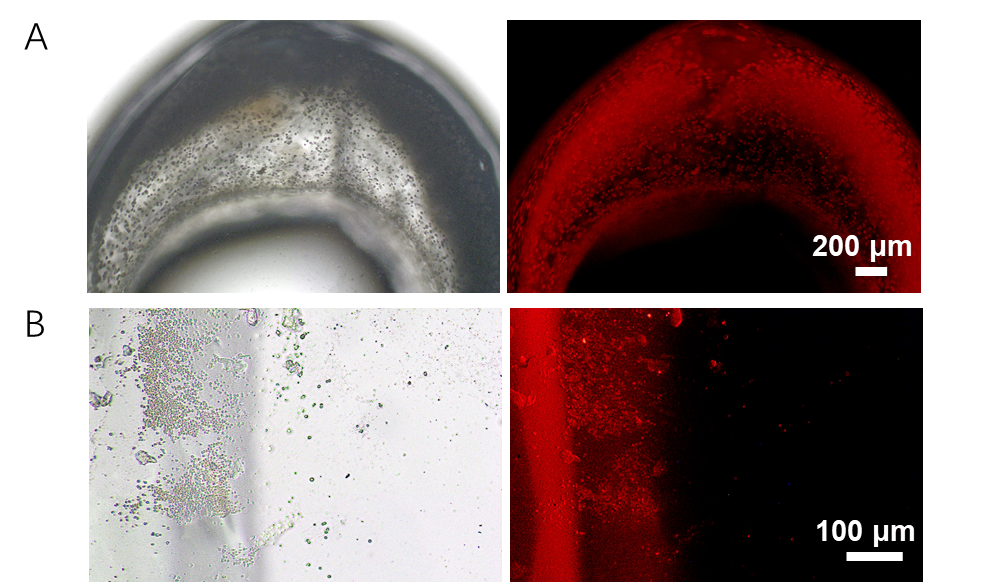

**Fig. S7. The growth of *C. vulgaris*.** (A) Bright field and fluorescent microscope images of *C. vulgaris*-laden SA/F127-DA/Ca^2+^ fibers, where *C. vulgaris* was immobilized in the shell phase and blank F127-DA in the core phase. (B) Proliferation of *C. vulgaris* immobilized in the shell phase into the F127-DA blank core phase hydrogel after 14 days of culture.
